# PepPr: De Novo peptide sequencing with peptide priors

**DOI:** 10.64898/2026.10.27.762126

**Authors:** Samuel Khan, Daniela Klaproth-Andrade, Yanik Bruns, Mathias Wilhelm, Julien Gagneur

## Abstract

Mass spectrometry-based peptide sequencing underpins proteomics. Database search confines to predefined candidates, whereas de novo peptide sequencing (DNPS) struggles with incomplete spectra. Existing hybrid methods use databases only for post hoc correction or retrieve previously identified spectra to guide decoding. We instead combine DNPS with a peptide prior: a language model fitted to a reference proteome, tolerating variants yet uncertain on strongly divergent sequences. Relative to prior-free DNPS, our implementation, PepPr, improved reference peptide identification and improved rather than suppressed variant identification. Using a related proteome as prior, it matched or outperformed homology-driven database search, and its gain over prior-free DNPS grew with relatedness. In antibody sequencing, it improved recovery of peptides spanning the variable-constant boundary, deamidation calls, and assembly, while preserving recall of hypervariable regions inaccessible to database search. Peptide priors thus bring reference knowledge into DNPS without limiting discovery.

## Introduction

Bottom-up tandem mass spectrometry is the most widely used analytical platform in proteomics. It captures the mass-to-charge ratios of ions produced by peptide fragmentation to generate a spectrum which, in principle, identifies its amino-acid sequence. However, peptide fragmentation is often incomplete, leaving parts of the peptide sequence ambiguous from the measurements alone^1^. It has long been recognized that database search, which leverages a reference proteome, helps circumvent the inherent ambiguity in experimental spectra^2,3^. For this purpose, database search evaluates a predefined set of candidate peptides, typically generated by in silico enzymatic digestion of the reference proteome of the species of interest. A peptide is then considered identified when its theoretical mass-to-charge fragments agree substantially with the observed tandem mass spectrum^4^. By assigning the complete sequence of the identified peptide, database search effectively imputes from the reference proteome amino acids at positions that are not directly resolved by the experimental fragment evidence.

The strength of database search is also its principal limitation: peptides absent from the candidate database cannot be identified. New database-search features have expanded that search space. These include increased mass tolerance^5^ and open search^1,6^ to capture post-translational modifications and amino acid variants, enzyme-unconstrained searches admitting peptides from non-canonical cleavage^7^, and variant-aware proteogenomic searches^7^. Still, the candidate set remains an explicit list of sequences chosen before the search begins.

For many applications, however, the relevant candidate set cannot be specified exhaustively in advance. This includes, but is not limited to, variants beyond single amino acid variants, including variants causing rare diseases, oncogenic and neoantigen-generating mutations^8^, variants of rapidly evolving viruses, and variants resulting from RNA editing, protein editing, or unannotated splicing. A further medically and biotechnologically relevant use case is antibody sequencing, where V(D)J recombination and somatic hypermutation expand the search space further^9^. Thus, even when the organism or sequence family is known, the exhaustive candidate-peptide listing that underpins database search can become prohibitive when the peptides of interest deviate from the reference.

De novo peptide sequencing aims to reconstruct peptide sequences directly from the spectral evidence. Modern de novo sequencing methods use deep learning to exploit the full richness of tandem mass spectra (reviewed by ref. 24). Although these methods have achieved substantial gains in accuracy, they are limited by the fundamental ambiguity of incomplete spectral evidence and therefore lag behind database search in terms of peptide recall.

This leaves an unmet need between database search and spectrum-only de novo sequencing. Several recent hybrid approaches have begun to close this gap. ReNovo^10^ and SearchNovo^11^ retrieve peptide-spectrum matches (PSMs) from large PSM databases during decoding. PowerNovo^12^ uses a BERT model trained on a diverse set of PSMs to weed out biologically unlikely sequences. However, these methods all leverage general-purpose PSM databases without prior information about the sequences expected in the sample of interest. The seminal model SMSNet^13^ is, to our knowledge, the only method that uses sample-specific prior information at all. However, it applies that prior as a post-processing step to correct low-confidence residues. Consequently, reference information can correct a fixed number of uncertain positions in the final prediction, but cannot shape which sequence hypotheses are explored by the neural network.

Here, we formalize peptide sequencing as combining two sources of evidence during inference: a spectrum-based next-amino-acid prediction, and a peptide prior language model that provides next-amino-acid probabilities from a reference proteome to disambiguate incomplete fragmentation. A small learned head fuses the prior with the base de novo sequencing predictions during autoregressive inference. This allows reference-proteome context to be used when it is informative while allowing spectral evidence to determine predictions outside the prior. Across species, variants, and antibody datasets, the prior behaved as intended: it helped most when spectral evidence was ambiguous, its benefit scaled with how relevant the prior was to the sample, and it remained compatible with the open-search behavior that motivates de novo sequencing. It was possible to integrate peptide priors with two state-of-the-art de novo models, suggesting a general mechanism. Together, these results establish peptide priors as a principled approach to bring reference-proteome context into open de novo sequencing, improving peptide recall while preserving the ability to identify sequences outside the prior.

## Results

### Integrating peptide priors into de novo sequencing

Since autoregressive decoding is currently the dominant de novo architecture, we designed an architecture to integrate with such a model. We had two main goals for the integration: The integrated model should provide reference sequence context when the current prefix closely matches a reference sequence, yet it should not override spectral evidence that deviates from the reference sequence. In order to make it easy to use priors for different use cases and species, we used a modular architecture. The prior is implicitly encoded in the weights of a separate model, and then integrated with the base de novo model using a small fusion head. We called our approach PepPr after the novel component, the peptide prior.

The prior distribution was defined operationally by sampling reference peptides together with controlled sequence perturbations (Fig. 1a). Specifically, the reference proteome was first digested in silico. Training batches were obtained by sampling peptides from this reference-peptide set randomly. For each sampled reference peptide, a triplet was generated comprising the intact peptide, a newly sampled single-amino-acid variant (SAAV), and a newly sampled peptide consisting of a prefix shared with the reference peptide followed by a random suffix. We used random suffixes, rather than fully random peptides, since C-termini of tryptic peptides are not random.

**Figure 1.**
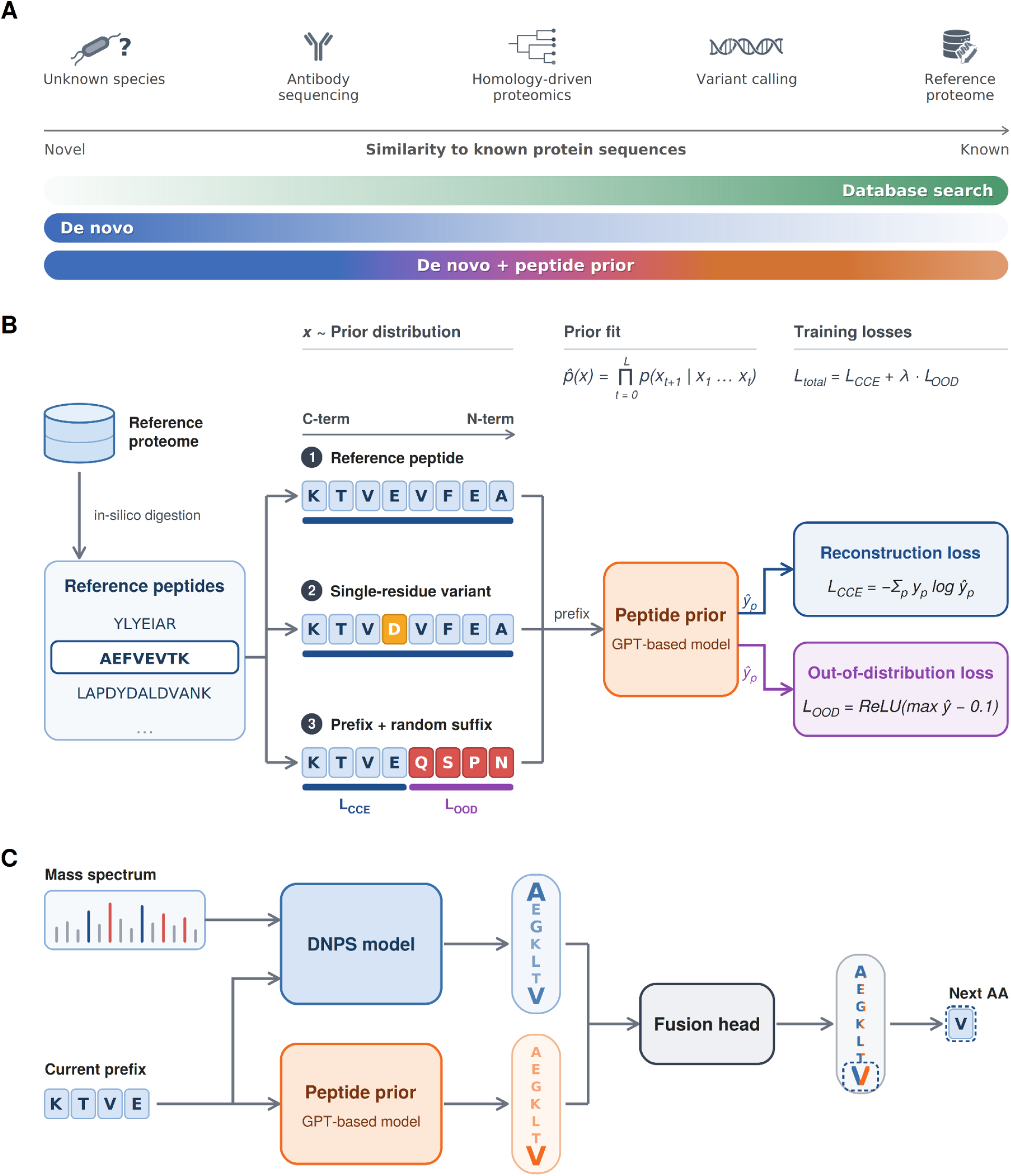
Integrating peptide priors into de novo sequencing. **a**, Peptide identification strategies across the range of sequence novelty. Proteomics applications investigated in this study are ordered by how similar peptides of interest are to known reference sequences, from unknown species to well-characterized reference proteomes. Database search performs best when the sample closely matches a reference proteome, whereas de novo sequencing does not rely on reference sequences and is therefore most valuable for novel ones. De novo sequencing with a peptide prior aims to span the full range: it draws on knowledge of known sequences where it applies while still being able to identify novel peptides. **b**, Peptide-prior training. Peptide prior models were trained to reconstruct in-silico digested reference peptides, read from the C- to the N-terminus. Training followed a scheme that sampled in equal parts: (1) unmodified reference peptides, (2) single amino acid variants of reference peptides, and (3) reference peptide prefixes joined to random suffixes. Predictions at positions in reference and variant sequences were optimized with a categorical cross-entropy reconstruction loss (L_CCE_). For random suffixes, an out-of-distribution loss (L_OOD_) penalized predicted probabilities above 0.1, encouraging uncertain predictions once the sequence diverged from the reference. The two losses were combined into a weighted sum. **c**, Prior-guided autoregressive decoding. At each decoding step, the base de novo sequencing model received the mass spectrum and the current sequence prefix, while the peptide prior received only the prefix. Their predictions were combined using a single-hidden-layer fusion head to assign probabilities to each possible next amino acid before beam search.

A transformer language model was trained by next-token prediction^14^ on these sequences to learn the prior distribution. No reference-proteome peptides were held out since the aim is to train a prior memorizing all reference peptides while allowing deviations to them. Each reference peptide was sampled on average 50 to 540 times during training, depending on proteome size. For simplicity, we refer to the resulting model as the peptide prior. For reference and SAAV sequences, the model was optimized to predict the next amino acid. For random-suffix sequences, we additionally applied an out-of-distribution (OOD) objective, that penalized high maximum softmax probabilities^15^ on positions within the random suffix, to avoid high-confidence predictions of the prior once the decoded sequence diverged from the reference-derived distribution.

At inference (Fig. 1b), the spectrum-conditioned de novo sequencing decoder and peptide prior each produced next-token logits for the current peptide prefix. A small fusion head takes as input both logit vectors and outputs a single logit vector for the next token prediction before beam search decoding (Methods). This way, the prior could supply sequence context while allowing spectral evidence to determine predictions outside the reference proteome. Because the fusion head takes the concatenated logits of the peptide prior and the de novo sequencing decoder and outputs a distribution over the decoder’s own vocabulary, the two models need not share a token set: the prior’s 20 canonical amino acids are combined with the decoder’s extended residue vocabulary, including modified residues. We trained a single fusion head per base model, which we reused across applications with application-specific peptide priors. The fusion head was trained on 200,000 reference-peptide PSMs from MassIVE-KB. To mitigate the fact that the training data consisted of reference peptides only, the prior’s top-1 and top-2 scores swapped during training for half of the tokens to simulate an incorrect prior. The fusion head therefore saw no real variant, splice, or non-reference peptides during training. Those were encountered only in the evaluations.

### Peptide priors improve identification of reference peptides over prior-free de novo sequencing

To assess the added value of PepPr, we used the nine species dataset, which is commonly used to assess de novo sequencing performance across different organisms and experimental setups^16^. For each species in this dataset, we integrated Casanovo^17^ with a peptide prior trained on the reference proteome of that species. The integration raised peptide recall by a median of +15.8 percentage points (pp), with improvements in all nine species (Fig. 2a, Supplementary Fig. 1). To put this performance improvement in context, we ran a suite of State of the Art de novo models on the mouse subset, confirming that PepPr beat the state of the art there (Fig 2b). Moreover, we showed that the method can transfer across autoregressive de novo architectures rather than being tied to one decoder, by integrating PepPr with ContraNovo^18^ as well, raising its average precision from 0.842 to 0.881 (Fig. 2b).

**Figure 2.**
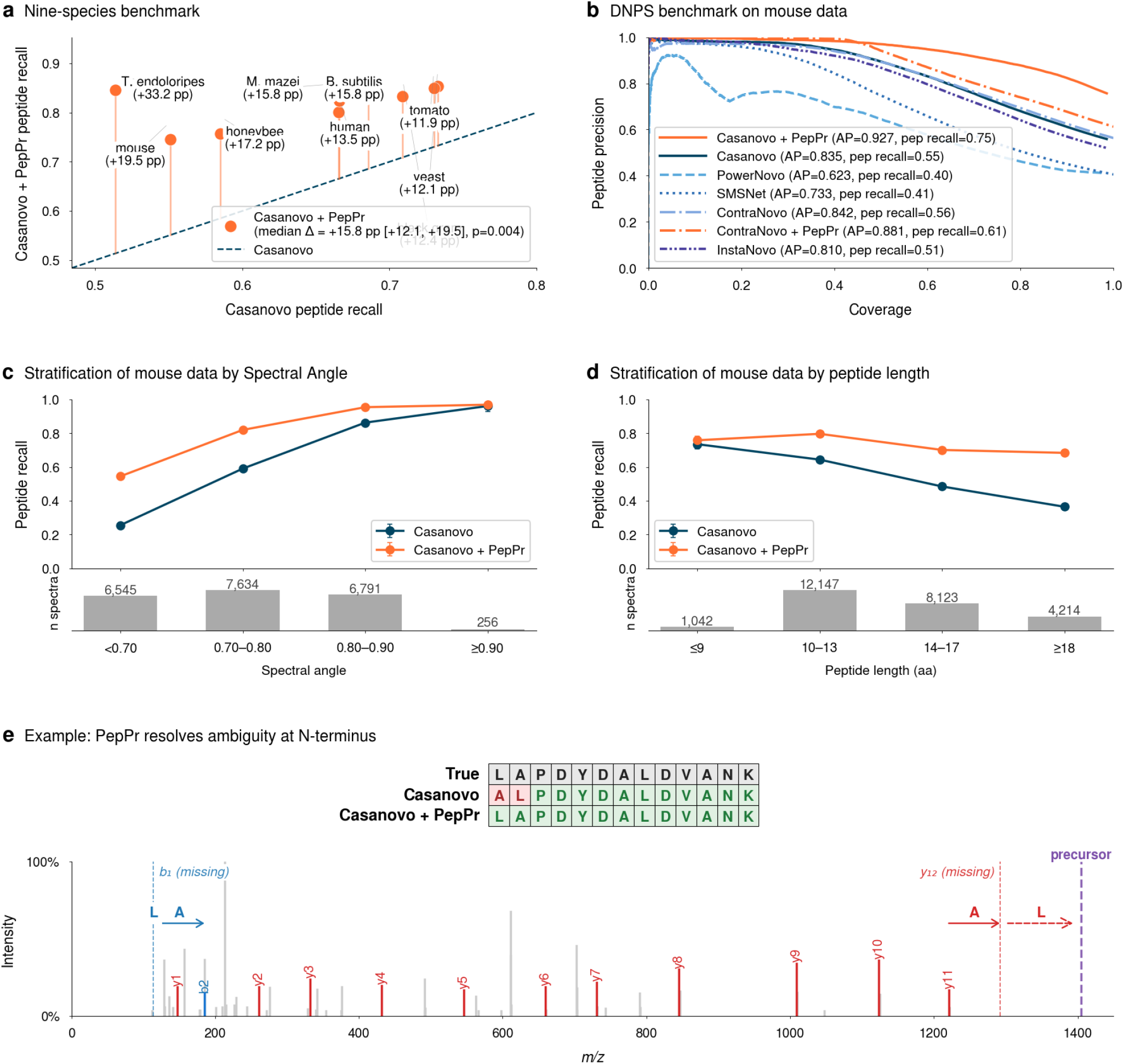
Peptide priors improve identification of reference peptides by resolving weak spectral evidence. **a**, Same-species peptide priors improve peptide recall across all nine species. Per-species peptide recall by Casanovo+PepPr using peptide priors trained on the tryptic digest of the reference proteomes of each respective species against the prior-free Casanovo baseline. Improvements (vertical distances to the diagonal) were found for all species (median +15.8 percentage points, *P* = 0.004, two-sided Wilcoxon signed-rank test), with the largest improvement of +33 percentage points on *T. endoloripes*. **b**, Precision against spectrum coverage when ranking peptide identification by each method’s confidence scores for the mouse dataset. Shown are the data for Casanovo+PepPr and ContraNovo+PepPr alongside their prior-free baselines, and the methods InstaNovo, SMSNet, and PowerNovo. **c,d**, Further stratification of the mouse data, showing peptide recall binned by **(c)** spectral angle (SA, computed against the Prosit-predicted spectrum of the ground-truth sequence) and **d,** length of the ground-truth sequence. **e**, Representative PSM from this data (scan 22758) illustrates the mechanism of improvement. The ground truth peptide was **<u>LA</u>**PDYDALDVANK. While prior-free Casanovo swapped the first N-terminal residues, Casanovo+PepPr correctly predicted the ground-truth ordering. The annotated spectrum marks the missing b1 and y12 peaks that make the correct order ambiguous from the spectrum alone.

We hypothesized that peptide priors help most when the spectrum itself provides the least information. To test this, we binned recall on the mouse subset by the spectral angle between the experimental spectrum and the Prosit^19^-predicted spectrum of the ground-truth sequence. Indeed, we observed the largest gains in recall on the lowest-quality spectra (Fig. 2c). As an additional check, we also binned recall by peptide length, since longer peptides are more likely to be missing peaks. Again, Casanovo’s recall fell from 0.74 for short peptides to 0.37 for long ones, whereas Casanovo+PepPr’s performance stayed at 0.68 or above across length buckets(Fig. 2d). Spectrum quality and peptide length also explained the difference in performance across species datasets. The largest improvement was observed for the *T. endoloripes* dataset, which had many low-spectral-angle spectra and the largest proportion of peptides consisting of 18 amino acids or more, whereas the mouse dataset combined a large number of shorter peptides with lower spectral quality and the *B. subtilis* dataset combined longer peptides with higher spectral quality (Supplementary Figs. 2 and 3).

Figure 2e shows a representative mouse PSM (scan 22758, true sequence **<u>LA</u>**PDYDALDVANK). Casanovo swapped the first two residues (**<u>AL</u>**PDYDALDVANK), whereas Casanovo+PepPr recovered the correct ordering. Even with high overall spectral agreement, the two singly-charged fragment ions that would fix this order, b_1_ and its complement y_12_, were both absent from the spectrum. This is a common ambiguity that spectral evidence alone cannot resolve because the first b-ions and the last y-ion are often missing^20^, . This example showcases the utility of a peptide prior: Up to two amino acids, the entire peptide sequence is supported by peaks. The prior helps fill the gaps, making the correct identification possible.

### Peptide priors improve, rather than suppress, single amino acid variant identification

An important deviation from the reference proteome database search assumption is caused by germline or somatic variants, notably in human genetics and in oncology^21^. Among these sequence alterations, single amino acid variants (SAAVs) represent the most frequent class of protein variation across individuals^22^ and tumors^23^. However, resolving SAAVs presents a well-known computational trade-off: expanded database search strategies face search-space and false-discovery penalties, while unconstrained *de novo* sequencing often lacks the sequence fidelity required to resolve flanking isobaric ambiguities^24^.

To evaluate whether PepPr resolves this trade-off, we first benchmarked Casanovo+PepPr on synthetic SAAV peptides from ProteomeTools^25^ (Fig. 3a). Each synthesized peptide in this dataset carries a single amino acid substitution relative to the canonical human reference proteome. Crucially, while PepPr’s prior was trained with randomly simulated substitutions, none of the specific ProteomeTools SAAV sequences were part of the reference peptidome nor seen during the training of the fusion head. Furthermore, because these spectra derive from pure synthetic pools searched against a restricted sequence library, the resulting ground-truth spectrum-peptide matches offer high confidence.

**Figure 3.**
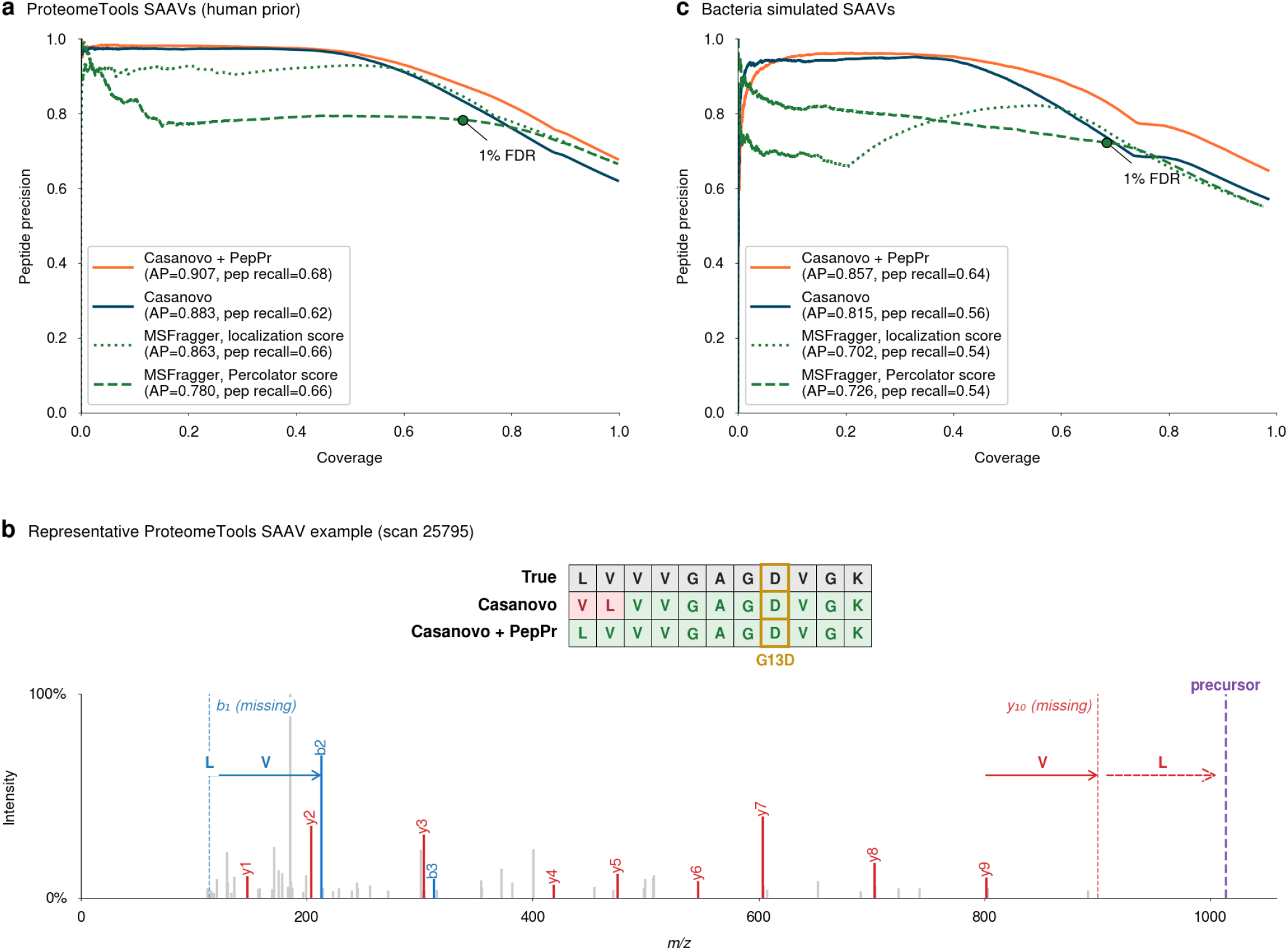
Peptide priors improve, rather than suppress, single amino acid variant identification. **a**, Precision-coverage curves on the ProteomeTools SAAV dataset, consisting only of peptides containing a single amino acid mutation from the reference proteome. Using a reference-proteome prior, Casanovo+PepPr achieves a peptide recall of 0.68 compared to Casanovo’s prior-free baseline of 0.62. The green curves plot database search baselines using the same reference proteome as the database and allowing for mass offsets corresponding to all possible single-amino-acid substitutions (MSFragger, ranked by Percolator score or by PTMProphet localization score). **b**, Representative ProteomeTools SAAV example (scan 25795). The table at the top shows the true peptide LVVVGAGDVGK which contains the KRAS G13D oncogenic mutation, the peptide predicted by Casanovo with a swap of the first two amino acids, and the correct predicted peptide by Casanovo+PepPr. The annotated spectrum below marks the missing b1 and y10 ions that would be required to resolve the ordering of the first two residues from spectral evidence alone. **c**, as in **a** for the Bacteria dataset.

In this benchmark, Casanovo+PepPr outperformed the prior-free Casanovo baseline, increasing peptide recall from 0.62 to 0.68 and average precision (AP) from 0.88 to 0.91 (Fig. 3a). To benchmark performance against leading database search engines, we also evaluated MSFragger^1^ in Detailed Mass Offset (DMO) mode^26^ using the canonical human reference proteome and all possible single-residue mass offsets. Casanovo+PepPr achieved comparable overall performance to MSFragger, whose AP ranged from 0.78 to 0.86 depending on the ranking metric (Methods). Across coverage thresholds, Casanovo+PepPr maintained higher precision (Fig. 3a). The selective correction of PepPr is exemplified by a peptide containing the oncogenic KRAS G13D mutation (wild type: LVVVGAGGVGK; variant: LVVVGAGDVGK), a frequent driver across colorectal cancers^27,28^ (Fig. 3b). Due to missing y_10_ and b_1_ peaks in the experimental spectrum, the ordering of the first two residues was ambiguous. While baseline Casanovo placed the G13D mutation correctly, it misassigned the N-terminus as a Leu/Val inversion (VLVVGAGDVGK). In contrast, Casanovo+PepPr recovered the true sequence (LVVVGAGDVGK). Here, the prior effectively resolved the flanking isobaric ambiguity toward the human reference backbone while preserving the spectrally supported variant residue, despite that specific variant being absent from the prior’s training data.

While ProteomeTools provide a strong ground truth, the spectra have a particularly high quality compared to spectra from actual biological samples. To assess the SAAV calling capacity in a more realistic setting, we borrowed the cross-species approach of Lundgren and colleagues^29^. Specifically, we analysed two samples of *Salmonella enterica* using the *E. coli* reference proteome, and assessed SAAV calling on the peptides differing by a single amino acid between the two species. On this dataset, Casanovo+PepPr with an *E. coli* prior outperformed both prior-free Casanovo and open database search both in average precision and in peptide recall (Fig. 3c).

Altogether, these results show that peptide priors do not trade variant discovery for reference-peptide recall. In both datasets, they improved recovery of non-reference substitutions over prior-free de novo sequencing and were competitive with open database search.

### Peptide priors substantially improve homology-driven proteomics

Proteomic studies of non-model organisms are often hampered by the lack of a well-assembled genome with high-quality protein annotation. Homology-driven proteomics circumvents this issue using the annotated proteome of a related species^30^. However, this approach breaks down as evolutionary distance increases and sequence divergence accumulates. Combining de novo sequencing with a related species’ prior could offer a flexible solution, but only if the prior boosts identification of conserved sequences without suppressing species-specific divergence.

To assess this potential benefit and its dependence on evolutionary distance, we evaluated PepPr with a human prior on the 14 non-human animal species of the Kingdoms of Life dataset^31^, and with an *E. coli* prior on ten bacterial species of the proteome resource of Abele *et al.*^32^. Peptide recall using closed database search decayed steeply as the exact tryptic overlap with the prior (Methods) decreased (Fig. 4a,b). Remarkably, prior-free Casanovo’s recall consistently and substantially outperformed homology-driven database search for all considered species pairs. Casanovo+PepPr matched prior-free Casanovo within 1.8 pp on the most distant species, and improved recall on closer ones by up to 10.5 pp (Fig. 4a,b). The gain increased with overlap in both kingdoms (Fig. 4c,d). Consistent observations were made on the nine species dataset (Supp Fig. 4).

**Figure 4.**
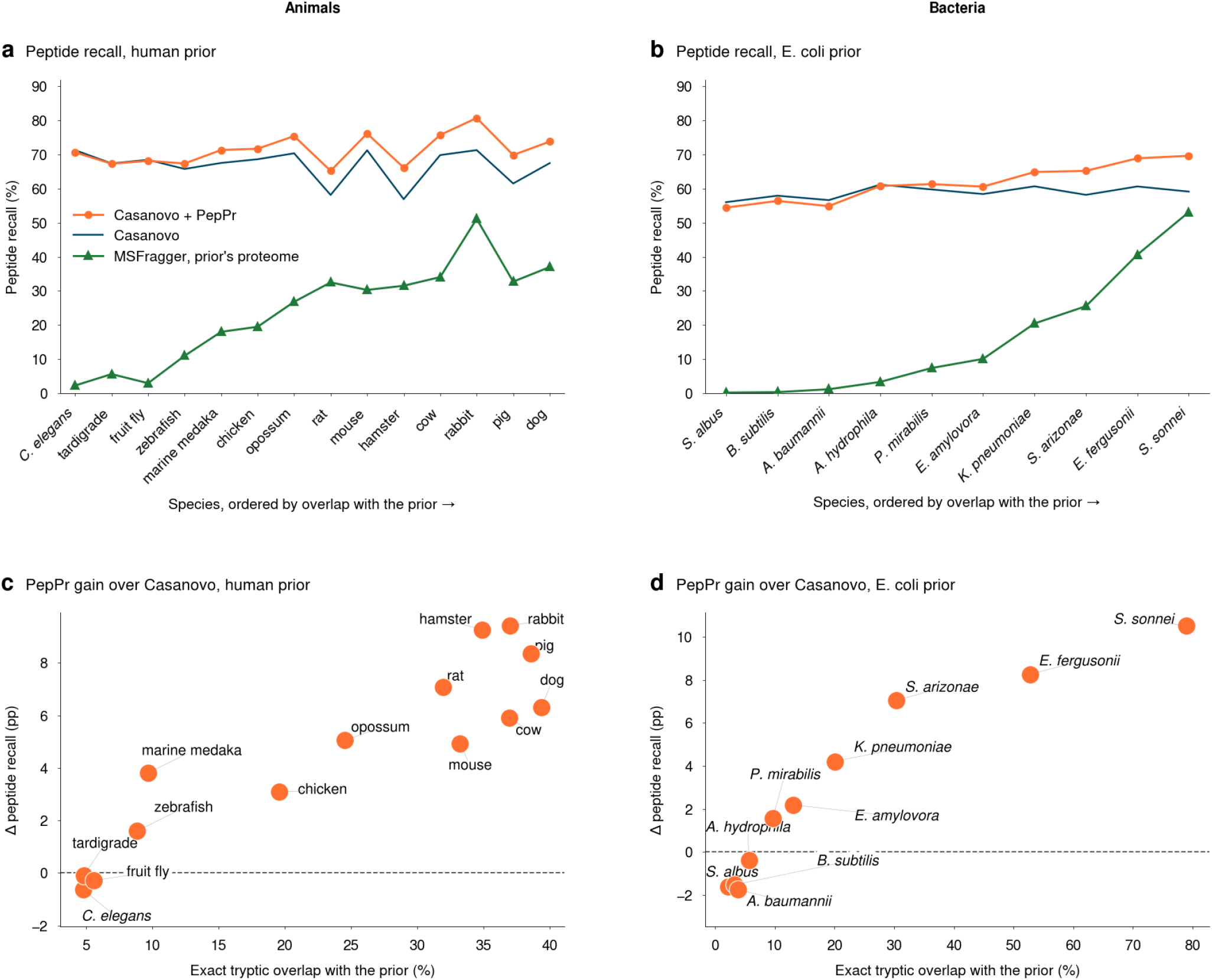
Peptide priors substantially improve homology-driven proteomics. a, PepPr improves peptide identification in homology-driven proteomics. Peptide recall of Casanovo+PepPr using a human prior (orange) on the 14 non-human species of the Kingdoms of Life dataset, with Casanovo (blue) and MSFragger (green) baselines. The MSFragger baseline was generated by searching over the same database that the PepPr Prior was trained on. Species are ordered by increasing exact tryptic overlap with the prior (Methods). **b,** as in **a** for the analogous experiment using an *E. coli* prior on other bacteria from the Bacteria dataset. **c,** PepPr benefits more from more informative cross-species priors. Improvement in peptide recall over prior-free Casanovo is plotted against the exact tryptic overlap with the prior (Methods). **d,** as in **c** for the Bacteria dataset.

Altogether, these results establish that PepPr provides a principled foundation for homology-driven proteomics. It exploits sequence conservation from model organisms to improve sensitivity, while leveraging the unconstrained ability of *de nov*o sequencing to discover lineage-specific sequences.

### Peptide priors improve antibodies sequencing

We next considered the problem of monoclonal-antibody sequencing, which are major therapeutic and research molecules^33^. Sequencing monoclonal antibodies is useful for molecular characterization, quality control, and bioengineering^34–36^. V(D)J recombination, junctional diversity, and somatic hypermutation create sample-specific sequence segments, making de novo sequencing the method of choice over database search^37^. On the other hand, framework and constant regions remain strongly constrained by germline and isotype sequence context. Therefore, we reasoned that monoclonal-antibody sequencing provides a useful test case for a peptide prior, which should use germline-derived information where it is informative and allow recovering diversified regions outside that prior. The importance of de novo sequencing in this domain is underscored by the recent introduction of XA-Novo^9^, a dedicated lab-workflow and assembly software designed specifically around antibody sequencing.

We evaluated six monoclonal antibodies: four from the XA-Novo dataset^9^ and the WIgG1 and IgG1 datasets distributed with Beslic *et al.*^39^. We excluded antibody mixtures from our analysis so that we could evaluate the downstream impact on assembly using the publicly available de novo antibody assembly software ALPS^40^. We aligned the peptides from these datasets to their published ground-truth reference sequences and categorized them by region. A peptide was called spanning if it overlapped with 1 residue or more in each of two adjacent regions. Peptides were hence categorized as CDR-only, framework-only (FR-only), constant-region-only (C-only), framework-CDR-spanning (FR*-*CDR), or variable/constant region-spanning (V-C). Casanovo+PepPr was run with peptide priors trained on mouse and human germline reference sequences digested without a tryptic assumption (Methods), and compared to the prior-free Casanovo baseline.

Casanovo’s baseline peptide recall was generally strong except on peptides spanning the V-C boundary (Fig. 5a). Because tryptic cleavage sites often lay at or near the V-C boundary, these fragments were primarily produced by non-tryptic proteases. Consequently, these non-tryptic peptides tended to be longer (Fig. 5b) and frequently lacked a basic C-terminal residue, potentially resulting in weaker y-ion ladders. PepPr dramatically improved recall within this category from 0.23 to 0.68. Crucially, this improvement did not come at the expense of pure CDR peptides, which were maintained or slightly improved. Recall improvements in the constant and framework regions were also observed. Consistent with the benefit of the peptide prior, germline-determined showed improved recall while performance on the somatically diversified CDR sequences were unaffected.

**Figure 5.**
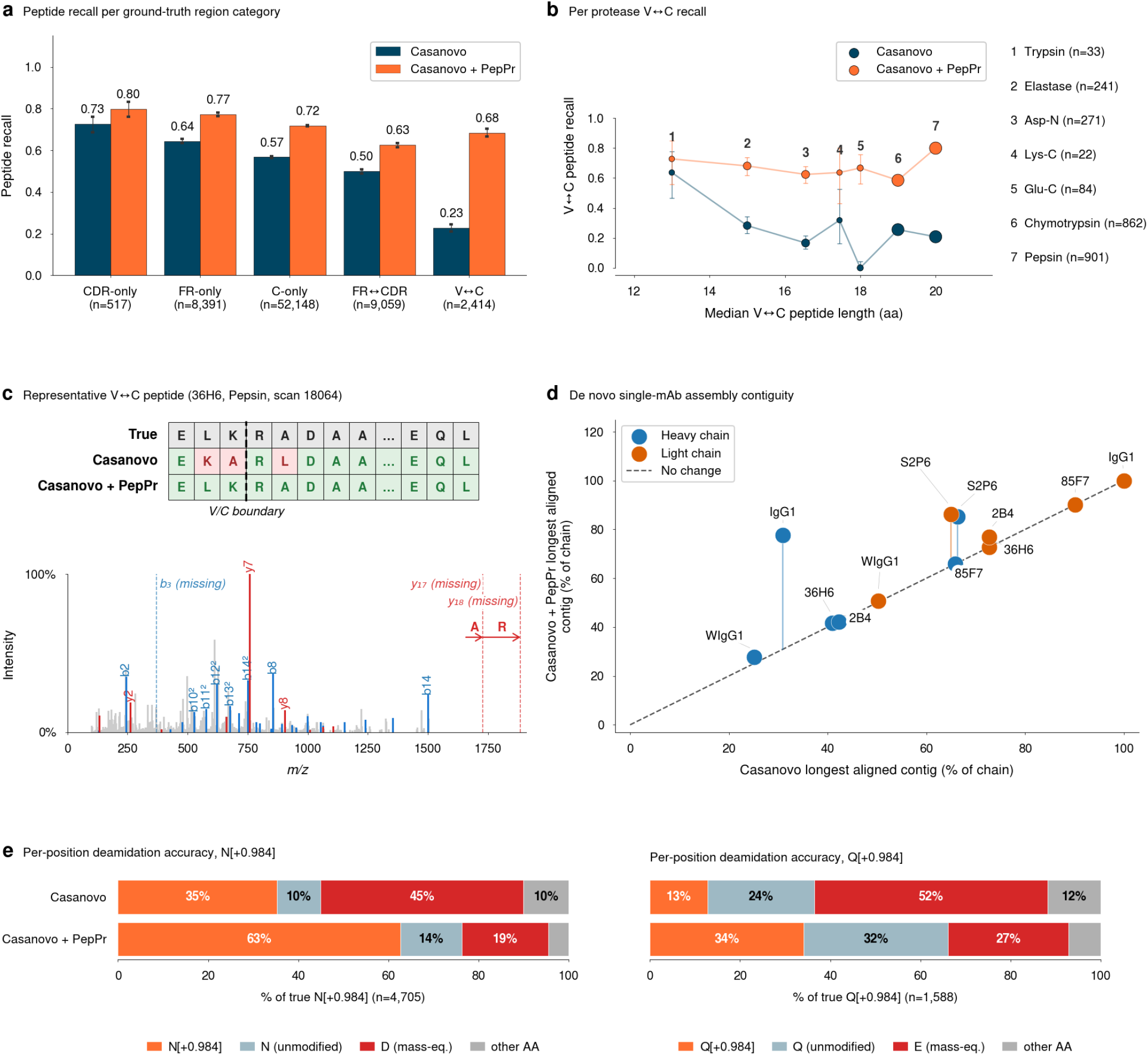
Peptide priors improve antibody sequencing. a, Peptide recall by region category: CDR-only, framework-only (FR-only), constant-region-only (C-only), framework-CDR-spanning (FR*-*CDR), or variable-constant region-spanning (V*-*C). Data were pooled across PSMs from six monoclonal antibodies: four from PXD060500 and two from MSV000079801. **b**, Recall of peptides spanning the V-C junction against peptide length per protease for Casanovo and Casanovo+PepPr. Bubble area is proportional to the number of spectra. The index table lists the proteases and the numbers of spectra. **c**, Representative peptide spanning the V*-*C junction (36H6 pepsin, scan 18064). Casanovo permutes amino acids at the positions 2 to 5, while Casanovo+PepPr matches the reference ordering. The annotated spectrum marks *b*_3_, *y*_17_ and *y*_18_ as missing. **d,** PepPr improves antibody assembly contiguity. For each antibody heavy (blue) and light (orange) chain, the longest aligned contig is shown as a fraction of the full reference-chain length for Casanovo+PepPr. The dashed diagonal represents Casanovo’s prior-free baseline. **e,** Accuracy of deamidation calls on asparagine (left) and glutamine (right) residues. At ground truth deamidated sites, predictions are classified as correct deamidation (dark blue), unmodified yet correct residue (light blue), mass-equivalent substitution (red, aspartate for asparagine and glutamate for glutamine), or any other amino acid (grey).

A representative V-C example from the pepsin digest of 36H6 illustrated the boundary-peptide effect (Fig. 5c). The true sequence, E**<u>LKRA</u>**DAA…EQL, contains an adjacent K–R pair, which is unexpected in tryptic training peptides because trypsin cleaves after both K and R. Casanovo returned E**<u>KARL</u>**DAA…EQL, permuting the boundary residues in a mass-preserving L/K/A 3-cycle at positions 2–5, whereas Casanovo+PepPr returned E**<u>LKRA</u>**DAA…EQL, recovering the canonical ordering.

We ran the ALPS assembler^40^ to test whether the peptide-level gains in peptide recall translated to improvements in downstream assembly metrics. In this analysis, Casanovo+PepPr produced longer aligned contigs (Fig. 5d).

The prior also improved deamidation calling, which is difficult for de novo tools because deamidated asparagine is mass-equivalent to aspartate, and deamidated glutamine is mass-equivalent to glutamate. Deamidation calling is also particularly relevant for antibody sequencing since long incubation times are often needed for multi-protease workflows, leading to high deamidation rates^34^. On true deamidation sites, PepPr raised correct recovery on asparagine and glutamine, chiefly by cutting the mass-equivalent mis-substitution rate for asparagine (Fig. 5e).

Together, these results show that the inclusion of peptide priors can be instrumental for antibody sequencing, where germline context can guide de novo sequencing without suppressing somatically diversified regions.

## Discussion

Here we introduced PepPr, a statistical framework for hybrid peptide sequencing that bridges the accuracy gap between de novo sequencing, which is limited by ambiguous spectra^24^, and database search, which cannot identify peptides outside its candidate list^1,4–7^, while preserving the capacity to detect peptides of unforeseen sequence. Earlier hybrid methods bring the database in through final-prediction correction or retrieval of general-purpose examples^10,11,13,41^, whereas here the prior acts during decoding, on the next-residue distribution. Our mechanism is simple and general: a peptide language model, trained on a reference proteome with explicit objectives for mutation tolerance and low out-of-distribution confidence, was fused with the per-step output of an autoregressive de novo peptide sequencing decoder. With such a prior, PepPr substantially improved median peptide recall on reference peptides across nine species^16^, improved rather than suppressed the identification of single amino acid variants, outperformed closed database search in homology-driven proteomics^30^, and substantially raised recall of antibody peptides spanning the variable-constant boundary while preserving recall of the hypervariable regions. Together, these results establish peptide priors as a principled way to use reference knowledge while preserving open discovery.

Viewed statistically, database search is a hard-lookup prior, uniform over a candidate list and zero elsewhere, and spectrum-only de novo sequencing is the case with no prior. A fitted prior can instead be fuzzy, tolerate variants, and yield to the spectrum on out-of-distribution sequences. PepPr therefore performs well on applications ranging from database-search-like applications, where most underlying peptides match the reference, to de novo applications, where it does not, and it is most useful in between. Because standard benchmarks such as the nine species dataset^16^ use closed database search as ground truth, the recall gains on reference peptides indicate that we recover much of what database search’s prior knowledge buys, without inheriting its hard constraint. We hope this will broaden the use of de novo sequencing in mass spectrometry, which remains heavily database-search based, since users no longer have to commit to a hard candidate list to benefit from prior knowledge. The framework is not tied to autoregressive decoding. A prior is a smaller model fitted to the distribution defined by the sampling process, and the same construction could be paired with a non-autoregressive decoder such as π-PrimeNovo^47^, which can use full bidirectional context to correct early mistakes. The integration would differ in implementation, and recent models that combine autoregressive and bidirectional representations^48^ suggest it is feasible, but the principle stays the same.

Several limitations remain. The applications we examined (reference peptides, variants, homology-driven proteomics, and antibodies) cover only part of where a peptide prior could help. Our evaluation focused on data-dependent acquisition of relatively high-quality spectra from high-resolution Orbitrap instruments. Settings with more ambiguous spectra, such as data-independent acquisition or single-cell proteomics, remain to be tested but may benefit even more from a peptide prior. The prior we investigated covers three modes: reference peptides, single-residue variants, and divergent sequences. Other sources of deviation, such as splicing, insertions, and deletions, were not modelled explicitly, and the relative proportions of the three components were not tuned. We deliberately used a single all-rounder prior to avoid specializing it to individual applications. This worked well empirically but may be suboptimal for a given use case. It can be a related proteome, as for venoms from non-model organisms^7,42^ and plants without annotated genomes^30,43^, where sample provenance is only approximately known. It can be a germline or host reference, as for antibodies^9,37^, neoantigens^8^ and rare-disease variants. Or it can be a partial or degraded reference, as in metaproteomics^44^ and paleoproteomics^45^. In PTM-aware models such as Modanovo^46^, a prior could help recall the base sequence and let the model concentrate on modification localization.

Furthermore, de novo sequencing still lacks established methodologies to assess and calibrate the confidence in their identifications independently of database-search ground truth, giving counterpart to the target-decoy-based false discovery rate^49^. Using peptide prior makes the problem even harder. Evaluation should move beyond database-search annotations, which favor reference peptides and so probably understate the benefit of non-reference sequences. Experimental data from synthetic peptides, such as ProteomeTools^25^, and consensus identifications across several search engines would give a more reliable benchmark. Another point concerns transparency. Imputing from prior knowledge underlies much of peptide identification, database search included, and is de facto needed, yet it is rarely traced or made visible to users. Hence, efforts to make the contribution of prior information transparent would be valuable, notably for medical diagnostic applications. While calibration, benchmarking, and transparency remain open and exciting lines of research, we foresee peptide priors being helpful across a wide range of mass spectrometry applications.

## Methods

### Datasets and reference sequences

#### MassIVE-KB

To train the fusion head, we used a 200,000-PSM subset of the Casanovo-published training set (10.5281/zenodo.14967861), randomly drawn from MassIVE-KB PSMs^17,53^. Peptide assignments from the Casanovo training set were used as ground-truth peptide assignments.

#### Nine species dataset

We used the revised dataset of Wen *et al.* ^16^ (MSV000090982), comprising tandem mass spectra from nine different species (between 25,526 and 1,358,337 PSMs each) for evaluation purposes in this study and to provide evaluation metrics comparable to other publications in the field. Peptide assignments from the existing database-search-derived annotations in the nine species dataset were used as ground-truth peptide assignments.

#### ProteomeTools SAAV

We considered the subset of ProteomeTools for model evaluation, which contained spectra of single–amino-acid-variant peptides (74,575 PSMs), each carrying exactly one single-residue substitution relative to a peptide from the human reference proteome. None of the variant sequences was included in the reference peptides of the prior. Peptide assignments from the MaxQuant search results published with the dataset were used as ground-truth peptide assignments.

#### Kingdoms of Life

We considered 14 non-human animal species of the Kingdoms of Life proteome resource of Müller *et al.* ^31^ (PXD014877), spanning *C. elegans* to rabbit. Peptide assignments from the MaxQuant search results as originally published (filtered by the authors at 1% PSM and protein FDR) were used as ground-truth peptide assignments. We restricted the dataset to 10,000 randomly chosen PSMs per species using reservoir sampling.

#### Bacterial dataset

To evaluate variant calling and homology-driven proteomics in bacteria, we used the bacterial proteome resource of Abele *et al.* ^32^ (MSV000096603). Peptide assignments from the MaxQuant searches published with the dataset were used as ground-truth peptide assignments. Because these searches were published without FDR filtering (PSM and protein FDR set to 100%), we applied a 1% PSM FDR ourselves by filtering to PSMs with a q-value ≤ 1%. For single-amino-acid variants, we followed Lundgren *et al.* ^29^ and treated *Salmonella* as a mutated *E. coli*. We used the 90-minute runs of *S. enterica* subsp. *arizonae* (DSM 9386) and *S. enterica* subsp. *diarizonae* (DSM 14847), comprising 89,275 PSMs, and classified each ground-truth peptide as reference, single-amino-acid variant, or other. For the homology analysis, we selected ten species of the dataset spanning 2% to 79% exact tryptic overlap with the *E. coli* prior and restricted each to 10,000 random PSMs of one 90-minute run.

#### Antibody datasets

We evaluated six single (not mixtures) monoclonal antibodies. Four came from the PRIDE entry PXD060500, introduced with the XA-Novo pipeline ^9^, each acquired after digestion by five proteases. Two additional antibodies, IgG1-Human and WIgG1-Mouse from MassIVE MSV000079801, were taken from the Beslic *et al.* dataset archive ^39,54^; their heavy and light chains were digested separately with other protease sets (Supplementary Table 3). Across the six antibodies, eight proteases were used in total. We did not use the Herceptin (trastuzumab) and anti-FLAG-M2 datasets from PRIDE PXD023419 ^38^ in the same archive, because their cysteines were alkylated with iodoacetic acid, while Casanovo only supports carbamidomethylated cysteine. To obtain peptide assignments, we ran MSFragger with Percolator rescoring via FragPipe (v23). Every digest was searched with the same settings: nonspecific cleavage, peptide lengths of 7 to 50 residues, 10 ppm precursor and 20 ppm fragment mass tolerance, isotope errors 0 to 3, carbamidomethylation of cysteine as a fixed modification, and oxidation of methionine and tryptophan and deamidation of asparagine and glutamine as variable modifications. The database contained the two reference chains published with each antibody and their reversed sequences as decoys, and PSMs were filtered at 1% FDR at the PSM and peptide level.

#### Reference proteomes

The peptide priors used for the nine species dataset, the Kingdoms of Life dataset, and the ProteomeTools dataset were trained on the canonical sequences of each species’ UniProt reference proteome (release 2026_01; reviewed Swiss-Prot and unreviewed TrEMBL entries), without isoforms except for the human proteome, which comprised Swiss-Prot entries only. The *E. coli* prior used for the bacterial data was trained for 200,000 iterations on the 4,399 Swiss-Prot entries of *E. coli* K-12 MG1655 (UP000000625) in the database published by Lundgren *et al.* ^29^. See Supplementary Table 1 for the exact proteome IDs and Supplementary Table 2 for the species name behind each label.

### Peptide prior training

#### Reference-proteome peptide sampling

Reference peptides were generated using an in silico tryptic digest (cleavage after every K and R) of each respective proteome, allowing for a maximum of one missed cleavage. The resulting sequences were deduplicated to form the final reference peptide prior set.

We defined the peptide prior operationally through a sampling procedure centered on these reference peptides. During training, reference peptides were sampled uniformly at random with replacement. Each training batch element consisted of a triplet derived from one reference peptide: (1) the reference peptide itself, (2) a single-residue variant obtained by substituting a random position with a different amino acid, and (3) a randomized-suffix sequence formed by concatenating a random-length prefix of the reference peptide with a random suffix.

#### Antibody peptide sampling

For the antibody peptide priors, construction differed in two aspects: the reference sequences and the *in silico* digestion method. Rather than utilizing an entire species’ proteome, the antibody priors sampled exclusively from immunoglobulin heavy and light chains. These sequences were computationally assembled using the enumerated V, C, and J alleles from IMGT/GENE-DB^55^. For the heavy chain, each V allele was generated as an independent FASTA sequence, alongside separate sequences representing every possible concatenation of J and C alleles. By generating these two distinct entry types for the heavy chain (and omitting the D region), we avoided modeling the extreme junctional diversity that causes the CDR3 region to deviate heavily from germline sequences. For the light chain, all possible direct concatenations of V, J, and C alleles were generated as independent entries. The resulting custom FASTA database contained 3,700 sequences for mouse and 16,900 for human. To accommodate the various proteases utilized in antibody sequencing experiments, the in silico digestion employed a protease-agnostic sliding window approach. Theoretical peptides were generated from the FASTA sequences by enumerating every L-mer, with L ranging from 6 to 25.

#### Hyperparameters and loss

The peptide prior is a decoder-only transformer with rotary position embeddings, sized like GPT-2 small: *L* = 12 layers, *d*_model_ = 768, *n*_heads_ = 12, FFN width 3072, and a vocabulary covering the 20 canonical amino acids, start and stop tokens, and an end-of-protein token. Peptides were presented in the C-to-N direction, matching Casanovo’s decoding direction. The prior’s next-token prediction inline] was trained with the standard next-token cross-entropy loss on the tokens of reference and variant sequences including the substituted residue. For tokens in the randomized suffix *S*, we instead penalized overconfidence with a bounded maximum-softmax penalty:

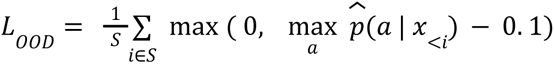

Models were optimized with AdamW^56^, a learning rate of 2 × 10*^−^*^4^ with linear warmup over 50,000 steps, cosine decay over the entire training duration, a batch size of 384 sequences (128 sampled reference peptides, each in the three versions described above) and a maximum of 750,000 training steps.

### Fusion with autoregressive de novo sequencing models

We trained a lightweight fusion head to combine the outputs of the de novo model with those from the peptide prior. For each base de novo model (Casanovo and ContraNovo), a fusion head was trained once on a subset of Casanovo’s training data and then reused unchanged with the various peptide priors. The fusion head’s input consisted of the concatenated next-token logits of the de novo model and the peptide prior model, and its output was a distribution over the base de novo vocabulary via one hidden layer of 128 units with ReLU activation. The de novo logits were normalized using batch normalization, which is appropriate because the backbone is fixed at inference time, so its running statistics remain valid, and the per-amino-acid calibration of its logits is preserved. The peptide prior logits were instead normalized using layer normalization, which standardizes each token’s logit vector independently and therefore introduces no running statistics tied to any particular prior.

Training minimized a cross-entropy loss against the reference token at each position, computed over individual token positions rather than whole peptides, with padding positions excluded. This gave roughly 4.5 million training tokens for the Casanovo head and 4.7 million for the ContraNovo head. We optimized with AdamW at an initial learning rate of 0.002 and a batch size of 8,192 tokens for 8 epochs, decaying the learning rate by a factor of 0.7 after each epoch.

Since the training data for the fusion head consisted solely of reference peptides, which correspond to cases where the peptide prior’s prediction matches the ground truth, we simulated cases where the peptide prior’s prediction was incorrect. To produce such samples, we applied a perturbation to the peptide prior’s outputs during training. For a randomly selected half of its output tokens, we swapped the top-1 and top-2 scores before passing them to the fusion head, leaving the de novo logits and the labels untouched. We primarily used Casanovo v5 with the released weights as the backbone de novo model. Casanovo was run with its default parameters except for beam search, where we used 5 beams.

### De novo sequencing baselines

ContraNovo, InstaNovo, PowerNovo and SMSNet were run on the mouse subset of the nine species dataset with their released weights. ContraNovo was run from its GitHub repository (commit 503fdb3) with the released checkpoint and its default configuration, using 5 beams. ContraNovo+PepPr used the same checkpoint and configuration together with the ContraNovo fusion head. InstaNovo v1.1.4 was run with the instanovo-v1.1.0 model and 5 beams instead of its default greedy decoding. PowerNovo v1.0.17 was run with its default beam size of 5 and BERT-based rescoring, with peptide assembly and protein inference disabled. SMSNet was run from its GitHub repository (commit facfaf4) with the released model without phosphorylation (m-mod), configured for that model as described in its documentation, with its default beam width of 20 and with rescoring.

### Database Search baselines

The Database search baselines used FragPipe 23.0 with MSFragger 4.2, Percolator 3.7.1, and Philosopher 5.1.1, with fully tryptic cleavage and up to two missed cleavages. A 20 ppm precursor and fragment mass tolerance was used, carbamidomethylation of cysteine was set as a fixed modification, and oxidation of methionine and protein N-terminal acetylation as variable modifications. While the ground truth PSMs derived from the searches published with each dataset had a minimum peptide length of 7, we used a minimum peptide length of 6 for the baseline searches to match Casanovo’s default. To report recall at full coverage, as for the de novo models, we scored every spectrum’s top-ranked peptide without FDR filtering, using the same amino acid matching routine applied to the de novo predictions. The databases were the same proteomes that the respective peptide prior was trained on.

For the ProteomeTools variant data, the baseline was a mass offset search designed to recover single amino acid substitutions. Rather than an unrestricted open search, we supplied a curated list of delta masses corresponding to every single amino acid substitution, each restricted to the residues that can give rise to it, and enabled localization of the delta mass so that shifted fragment ions were included during scoring. Delta mass sites were localized with PTMProphet, falling back to the MSFragger localization score where PTMProphet returned no result. Because PTMProphet reads spectra from mzML, the mass-offset searches ran on all MS2 spectra of each run and were scored on the ground-truth spectra.

For the *Salmonella* variant dataset, we ran the same mass-offset search on the two 90-minute runs, against the *E. coli* K-12 MG1655 proteome of Lundgren *et al.*^29^.

For the Kingdoms of Life and bacterial homology-driven proteomics searches, the baseline instead was a conventional closed database search in which each species’ spectra was searched against the proteome the prior was trained on (human, or *E. coli*) rather than its own. By construction, the recall of these search results was bounded by the overlap.

### Evaluation

We reported peptide-level precision, peptide recall, and average precision, defined as the area under the precision–coverage curve when peptides are ranked by each model’s reported peptide-level score using scikit-learn’s *auc* function^68^. Following Casanovo’s definition, peptide coverage was computed against all spectra in the dataset. Spectra for which a method emitted no prediction (e.g., due to peak-count filtering) counted as not recovered, so no method was rewarded for not providing a prediction on low-quality spectra. A predicted residue counted as correct when its mass was within 0.1 Da of the reference residue and the cumulative prefix masses agreed within 0.5 Da, and a peptide counted as correct when all residues matched. This made the metrics insensitive to mass-equivalent ambiguities such as isoleucine/leucine but also equated other residue pairs within 0.1 Da, such as deamidated asparagine and aspartate. Peptide recall is the fraction of all spectra for which the predicted peptide was correct under this rule. Since spectra without a prediction are counted, peptide recall equals the precision at the end of the precision-coverage curve. Deamidation calls (Fig. 5e) were scored by residue identity instead .

For spectral angle stratification, we predicted the b/y fragment intensities of each ground truth peptide with Prosit ^19^, accessed through KoinaPy/Koina ^57^. After matching the predicted spectrum R to the observed spectrum I, spectral angle was computed as 1 *−* (2*/π*) arccos(cos_sim(_*^√^I, ^√^R*)), where cos_sim_ is the cosine similarity. The resulting values range from 0 to 1, with 1 indicating the highest similarity between the observed spectrum and the Prosit predicted ground-truth.

For the homology analyses, we quantified how much of a target species’ peptidome a prior covers by its exact tryptic overlap. We define this as the percentage of peptides obtained by digesting the species’ proteome in silico that occur exactly in the prior’s training set, with isoleucine and leucine treated as identical.

For antibody-region analyses, each antibody peptide was aligned to its reference sequence and assigned to a region category: CDR-only, framework-only (FR-only), constant region-only (C-only), peptides spanning a framework-CDR boundary (FR*-*CDR), and peptides spanning the variable-constant boundary (V*-*C). Region-category recall was pooled across all antibody and protease combinations and reported with Wilson 95% confidence intervals. For the per-protease V-C boundary analysis, we pooled PSMs across all six single antibodies for each protease and computed peptide-level recall for Casanovo and Casanovo+PepPr, together with the median V*-*C peptide length. Region boundary positions came from IMGT.

For antibody assembly, per-scan peptide predictions were pooled across proteases for each antibody and arm, filtered to predictions with peptide-level score *≥* 0.1 to exclude known-bad predictions, and assembled with ALPS^40^. We ran ALPS with the top *c* = 20 contigs for each *k* from 7 to 11, then unioned and deduplicated the resulting per-*k* contig sets. Each contig (length typically 80-200 AAs) was aligned to the mature heavy- and light-chain antibody reference with npysearch^58^ (BLOSUM62 seed-and-extend, maxAccepts= 5). High-scoring segment pairs (HSPs), the local alignments between a contig and the reference, with *>* 95% identity were accepted for all assembly metrics. Accepted HSPs were used to compute reference coverage and longest single-contig alignment. As an upper bound, we reported the full mature heavy- or light-chain reference length.

For deamidation recovery, we pooled all ground-truth deamidated asparagine and glutamine sites across the six antibodies and their proteases. Unlike the peptide-level metrics, this analysis compared residues by identity rather than by mass, since deamidated asparagine and glutamine are mass-equivalent to aspartate and glutamate. We used the spectra predicted by both Casanovo and Casanovo+PepPr for which both predictions had the same number of residues as the ground-truth peptide, so that positions could be compared directly. At each ground-truth deamidated position, the predicted residue at the same position was classified as the deamidated residue (correct), the unmodified residue (missed modification), the mass-equivalent native residue created by the +0.984 shift (aspartate or glutamate; mis-substituted), or any other residue, including other modifications.

### Statistics and reproducibility

The same-species and cross-species paired median Δ peptide-recall values are each reported with a bootstrap 95% confidence interval on the median and a two-sided Wilcoxon signed-rank *P-*value. The Kingdoms of Life overlap-versus-lift correlation is a Pearson *r* over the 26 pooled (species *×* prior) points. The assembly-contiguity and deamidation summaries and the single-PSM worked examples are point estimates without confidence intervals.

Models are implemented in PyTorch 2.7.0^59^. The code builds on Casanovo 5.1 (forked to add the fusion head) and depthcharge 0.4.9^17^, and nanoGPT^60^ for peptide priors. Mass-spectrometry peak files are parsed with pyteomics 4.7.5^61,62^, and annotated tandem mass spectra are rendered with spectrum_utils 0.5.0^63^.

De novo contigs for assembly analysis are built with ALPS^40^ and aligned with npysearch 1.3.1^58^; FASTA/sequence I/O uses Biopython 1.85^64^. Numerical analysis uses NumPy 1.26.4^65^ and pandas 2.2.3^66^; statistics use SciPy 1.15.3^67^; precision–coverage areas use scikit-learn 1.7.0^68^.

## Acknowledgements

We would like to thank the members of the Gagneur and Wilhelm labs for their valuable feedback and fruitful discussions, especially Joel Lapin, Wassim Gabriel, Thomas Zoeschg, Niklas Feurstein, Kurt Stonus, Ayla Schröder, Jesse Angelis, Petra Navarcikova, Qussai Abbas, Sonja Stockhaus, and Christian Nix. We are also grateful to members of the Kuster and Wilhelm labs for their support with data transfer, and to the Casanovo team for their help in reproducing their published results.

## Author contributions

J.G. conceived the original idea for the method, which S.K. developed into the approach presented here. S.K. implemented the peptide prior and the fusion network, ran the experiments, and wrote the manuscript. D.K.-A. and Y.B. contributed to method development, dataset curation, and evaluation. D.K.-A., M.W. and J.G. supervised the work. All authors contributed to interpretation of the results and revision of the manuscript.

## Funding and additional information

This study was supported by the German Bundesministerium für Bildung und Forschung (BMBF) supported through the project CLINSPECT-M [16LW0243 K to Y. B., D. K., J. G.]; by the Deutsche Forschungsgemeinschaft (DFG, German Research Foundation) via the IT Infrastructure for Computational Molecular Medicine [#461264291, #553375143] and the collaborative research centers [TRR179-TP24, TRR237-A07.]; by the European Research Council via an ERC Starting Grant [#101077037 to MW] and by the State of Bavaria (BayVFP 2024–2027 “P3M“). Views and opinions expressed are those of the author(s) only and do not necessarily reflect those of the European Union or any other granting authority, which cannot be held responsible for them.

## Competing interests

The authors declare the following financial interests/personal relationships which may be considered as potential competing interests: Mathias Wilhelm is a scientific advisor of OmicScouts GmbH, a Momentum Biotechnologies Company, with no operational role. The remaining authors declare no competing interests.

## Code and data availability

A reference implementation of the method is available at https://github.com/gagneurlab/PepPr, and the analysis scripts to reproduce all figures and results are available at https://github.com/gagneurlab/PepPr_paper. Pretrained model weights for the peptide priors and the fusion network, together with the FASTA files they were trained on, are available on Zenodo (https://doi.org/10.5281/zenodo.22252036). The processed benchmark spectra with their ground-truth annotations (nine-species, Kingdoms of Life subsets, ProteomeTools SAAV, antibody datasets and the fusion-head training subset of MassIVE-KB), together with all predictions, database-search baseline results and evaluation outputs, are available on Zenodo (https://doi.org/10.5281/zenodo.21869640; the files are accessible to peer reviewers through a private link and will be public upon publication). The raw data are available from the original repositories: nine-species (MSV000090982), Kingdoms of Life (PXD014877), antibodies (PXD060500, MSV000079801), bacterial resource (MSV000096603), the *E. coli* database of Lundgren *et al.*^29^ (MSV000092029), Casanovo training data (https://doi.org/10.5281/zenodo.14967861) and ProteomeTools SAAV (PXD081052, available to peer reviewers upon request).

## Supplementary Figures

**Supplementary Figure 1.**
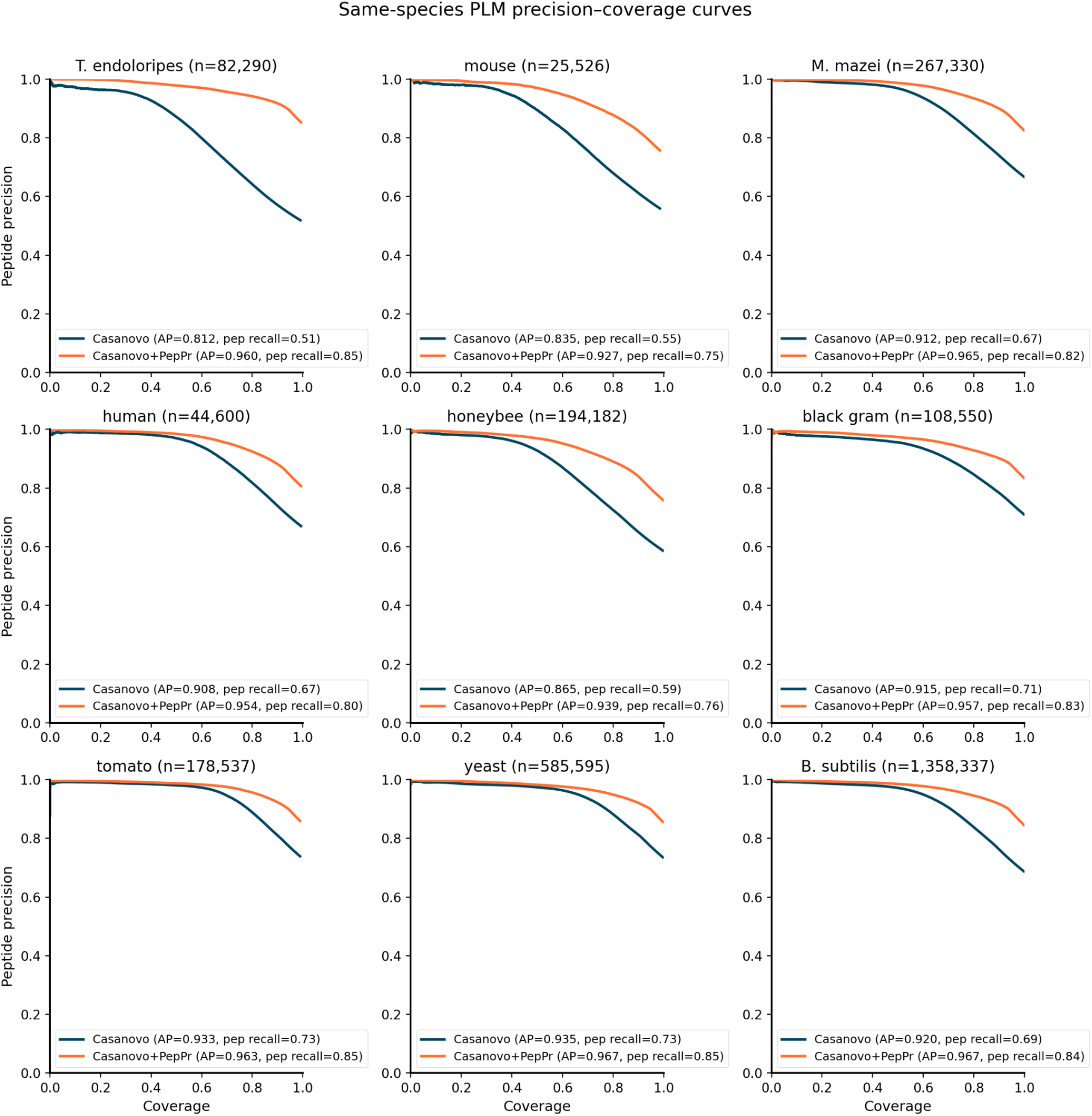
Per-species precision–coverage curves with same-species priors. Precision–coverage curves underlying the same-species scatter in Fig. 2a: for each of the nine species, Casanovo versus Casanovo+PepPr using a same-species prior (with the shared MassIVE-KB fusion head), with the average precision (AP) of each curve annotated. Casanovo+PepPr improves the AP in every species.

**Supplementary Figure 2.**
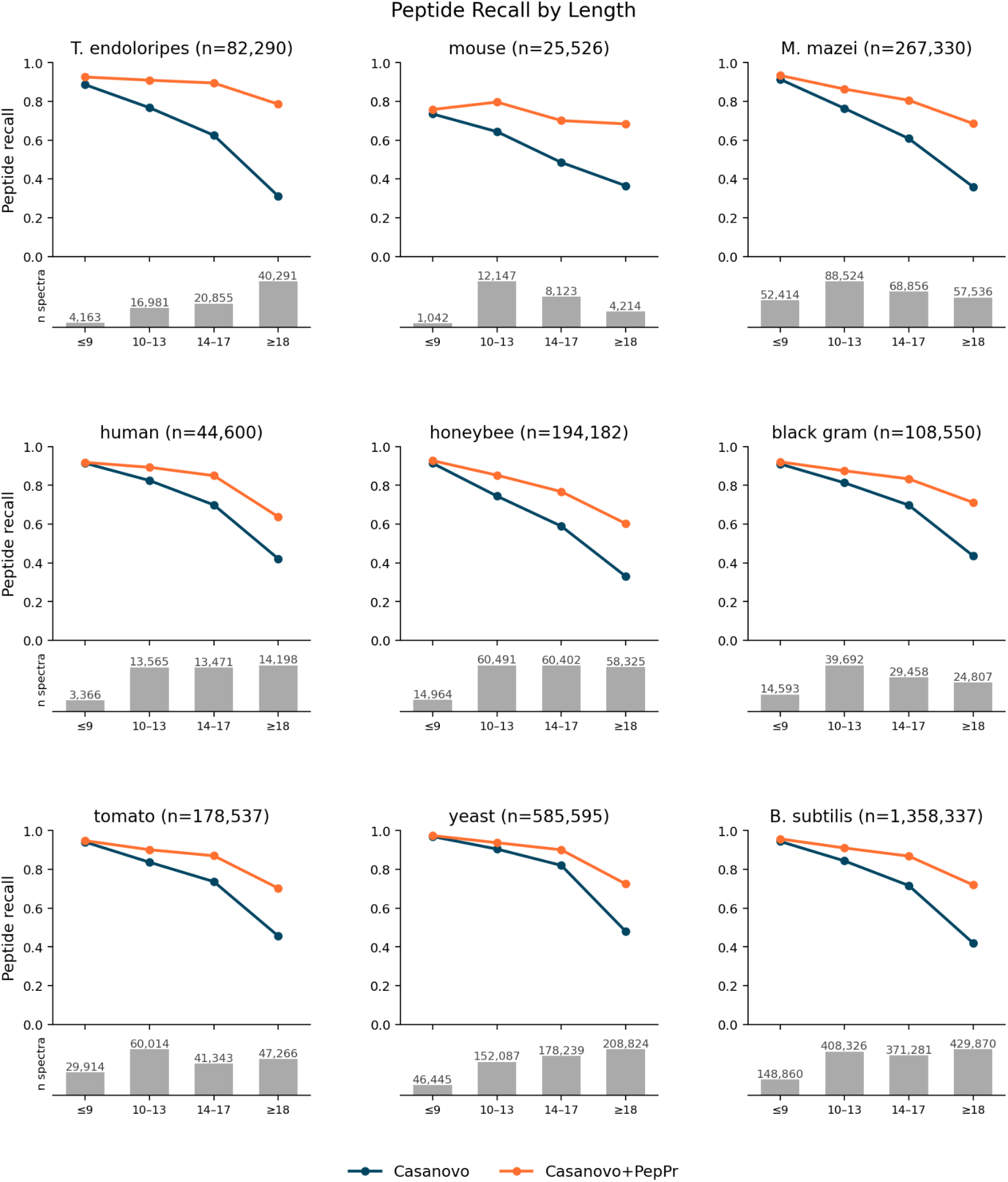
Peptide precision by length across all nine species. Per-species version of Fig. 2d: peptide recall for Casanovo and Casanovo+PepPr stratified by true peptide length (*≤* 9, 10–13, 14–17, *≥* 18 aa), with per-bin spectrum counts shown below each panel, for all nine species of the benchmark (cf. the mouse subset, Fig. 2d).

**Supplementary Figure 3.**
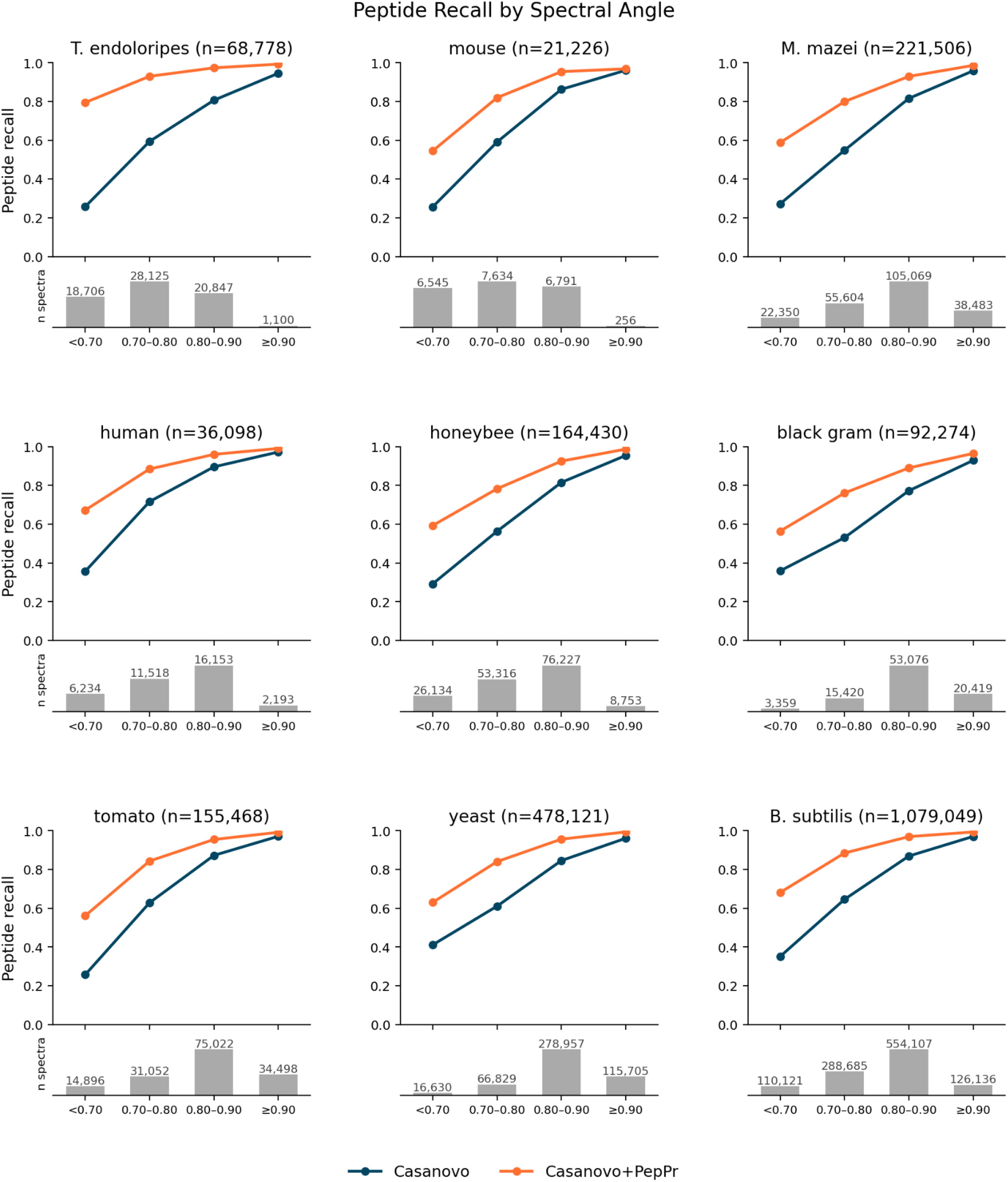
Peptide precision by spectral angle across all nine species. Per species version of Fig. 2c: peptide recall for Casanovo and Casanovo+PepPr stratified by spectral angle using fixed spectral-angle bins (*<* 0.70, 0.70–0.80, 0.80–0.90, *≥* 0.90). Spectra are the same species-level inputs as in Supplementary Fig. 2, except that spectra without a Prosit prediction are removed; this prediction-coverage filter accounts for the count differences between the two supplementary stratifications. Retained per-bin spectrum counts are shown below each panel. The largest Casanovo+PepPr versus Casanovo differences occur in the bins of lowest spectral angles.

**Supplementary Figure 4.**
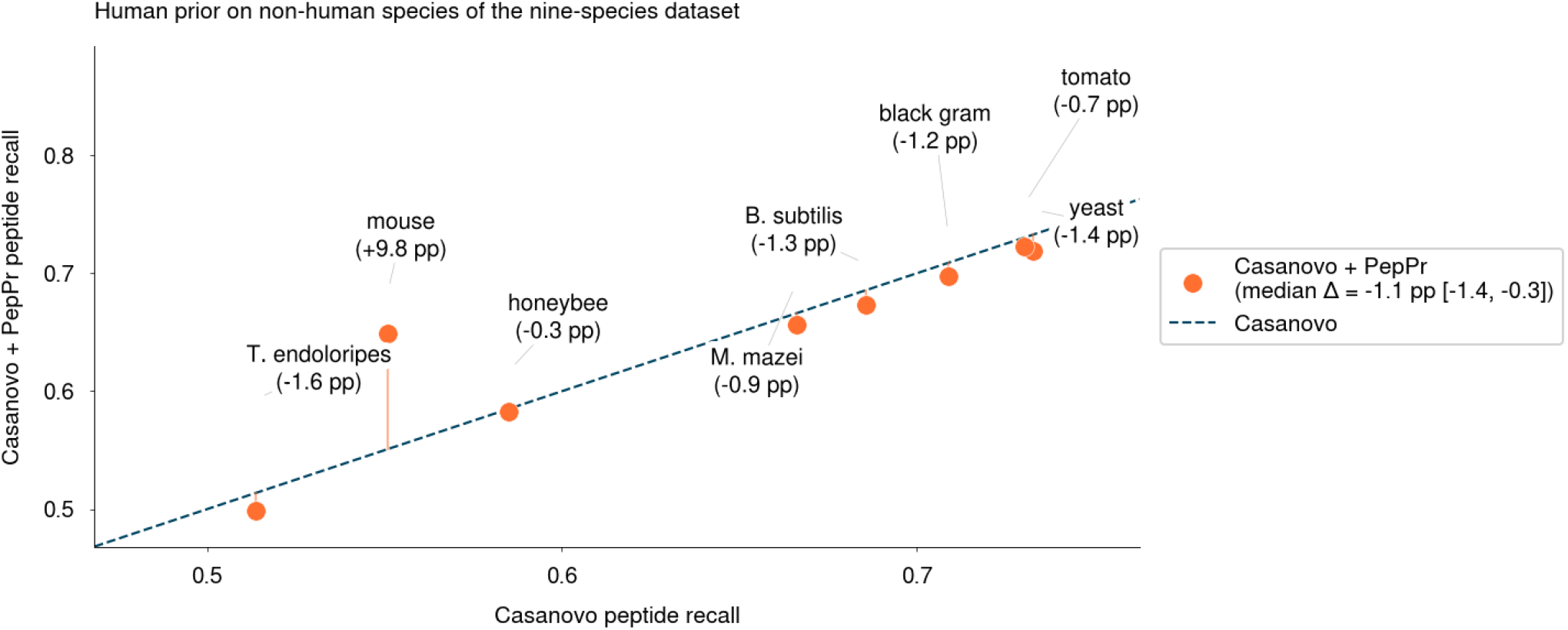
Peptide recall of Casanovo+PepPr using a human prior on the 8 non-human species of the nine species dataset. The diagonal represents Casanovo’s prior-free recall.

**Supplementary Table 1.**

| Species label | Proteome ID | UniProt version | Includes Isoforms | Swiss-Prot Entries | TrEMBL Entries |
| --- | --- | --- | --- | --- | --- |
| human | UP000005640 | 2026_01 | yes | 42,562 | 0 |
| mouse | UP000000589 | 2026_01 | no | 17,188 | 4,664 |
| Baker's yeast | UP000002311 | 2026_01 | no | 6,066 | 0 |
| *B. subtilis* | UP000001570 | 2026_01 | no | 4,191 | 80 |
| honeybee | UP000005203 | 2026_01 | no | 86 | 9,849 |
| tomato | UP000004994 | 2026_01 | no | 511 | 34,152 |
| black gram | UP001374535 | 2026_01 | no | 0 | 37,670 |
| *M. mazei* | UP000034578 | 2026_01 | no | 0 | 3,460 |
| *T. endoloripes* | UP000094849 | 2026_01 | no | 0 | 3,951 |
| *E. coli* | UP000000625 | 2022_01 | no | 4,399 | 0 |

**Supplementary Table 2.** Species labels used in figures and corresponding binomial names.

| Label in figures | Species |
| --- | --- |
| Human | *Homo sapiens* |
| Mouse | *Mus musculus* |
| Cow | *Bos taurus* |
| Rabbit | *Oryctolagus cuniculus* |
| Fruit fly | *Drosophila melanogaster* |
| C. elegans | *Caenorhabditis elegans* |
| Marine medaka | *Oryzias melastigma* |
| Black gram | *Vigna mungo* |
| Tardigrade | *Hypsibius dujardini* |
| Opossum | *Monodelphis domestica* |
| T. endoloripes | *Thiodiazotropha endoloripes* |
| B. subtilis | *Bacillus subtilis* |
| M. mazei | *Methanosarcina mazei* |
| Baker's yeast | *Saccharomyces cerevisiae* |
| Honeybee | *Apis mellifera* |
| Tomato | *Solanum lycopersicum* |
| Zebrafish | *Danio rerio* |
| Chicken | *Gallus gallus* |
| Rat | *Rattus norvegicus* |
| Hamster | *Cricetulus griseus* |
| Pig | *Sus scrofa* |
| Dog | *Canis lupus familiaris* |
| S. sonnei | *Shigella sonnei* DSM 5570 |
| E. fergusonii | *Escherichia fergusonii* DSM 13698 |
| S. arizonae | *Salmonella enterica* subsp. *arizonae* DSM 9386 |
| K. pneumoniae | *Klebsiella pneumoniae* DSM 30104 |
| E. amylovora | *Erwinia amylovora* DSM 30165 |
| P. mirabilis | *Proteus mirabilis* DSM 4479 |
| A. hydrophila | *Aeromonas hydrophila* DSM 30187 |
| A. baumannii | *Acinetobacter baumannii* DSM 30007 |
| S. albus | *Streptomyces albus* DSM 40313 |

**Supplementary Table 3.** Number of spectra per antibody and protease used in Fig. 5a,b.

| Antibody | Trypsin | Chymotrypsin | AspN | Elastase | Pepsin | GluC | LysC | Proteinase K |
| --- | --- | --- | --- | --- | --- | --- | --- | --- |
| 2B4 | 1,956 | 2,147 | 2,709 | 2,623 | 3,420 | – | – | – |
| 36H6 | 1,622 | 1,942 | 2,063 | 2,863 | 3,450 | – | – | – |
| 85F7 | 2,253 | 2,234 | 3,178 | 2,518 | 3,347 | – | – | – |
| S2P6 | 1,870 | 2,929 | 2,571 | 2,736 | 4,001 | – | – | – |
| IgG1-Human heavy chain | – | 2,247 | 1,419 | – | – | 2,077 | 1,920 | 1,854 |
| IgG1-Human light chain | – | 2,259 | 2,302 | – | – | 999 | 812 | 1,333 |
| WlgG1-Mouse heavy chain | – | 781 | 957 | – | – | – | – | – |
| WlgG1-Mouse light chain | – | 458 | 679 | – | – | – | – | – |
| Total | 7,701 | 14,997 | 15,878 | 10,740 | 14,218 | 3,076 | 2,732 | 3,187 |

## References

1. Kong, A. T., Leprevost, F. V., Avtonomov, D. M., Mellacheruvu, D. & Nesvizhskii, A. I. MSFragger: ultrafast and comprehensive peptide identification in mass spectrometry-based proteomics. Nature Methods 14, 513–520 (2017).

2. Eng, J. K., McCormack, A. L. & Yates, J. R. An approach to correlate tandem mass spectral data of peptides with amino acid sequences in a protein database. Journal of the American Society for Mass Spectrometry 5, 976–989 (1994).

3. Mann, M. & Wilm, M. Error-tolerant identification of peptides in sequence databases by peptide sequence tags. Analytical Chemistry 66, 4390–4399 (1994).

4. Steen, H. & Mann, M. The ABC’s (and XYZ’s) of peptide sequencing. Nature Reviews Molecular Cell Biology 5, 699–711 (2004).

5. Chick, J. M. et al. A mass-tolerant database search identifies a large proportion of unassigned spectra in shotgun proteomics as modified peptides. Nature Biotechnology 33, 743–749 (2015).

6. Yu, F. et al. Identification of modified peptides using localization-aware open search. Nature Communications 11, 4065 (2020).

7. Nesvizhskii, A. I. Proteogenomics: concepts, applications and computational strategies. Nature Methods 11, 1114–1125 (2014).

8. Bassani-Sternberg, M. & Coukos, G. Mass spectrometry-based antigen discovery for cancer immunotherapy. Current Opinion in Immunology 41, 9–17 (2016).

9. Xiong, Y. et al. XA-Novo: high-throughput mass spectrometry-based de novo sequencing technology for monoclonal antibodies and antibody mixtures. Nature Communications 17, 3391 (2026).

10. Chen, S., et al. ReNovo: retrieval-based de novo mass spectrometry peptide sequencing. In International Conference on Learning Representations (2025).

11. Xia, J., et al. Bridging the gap between database search and de novo peptide sequencing with SearchNovo. In International Conference on Learning Representations (2025).

12. Petrovskiy, D. V. et al. PowerNovo: de novo peptide sequencing via tandem mass spectrometry using an ensemble of transformer and BERT models. Scientific Reports 14, 15000 (2024).

13. Karunratanakul, K., Tang, H.-Y., Speicher, D. W., Chuangsuwanich, E. & Sriswasdi, S. Uncovering thousands of new peptides with sequence-mask-search hybrid de novo peptide sequencing framework. Molecular & Cellular Proteomics 18, 2478–2491 (2019).

14. Radford, A., Narasimhan, K., Salimans, T. & Sutskever, I. Improving language understanding by generative pre-training. OpenAI (2018).

15. Hendrycks, D. & Gimpel, K. A baseline for detecting misclassified and out-of-distribution examples in neural networks. In International Conference on Learning Representations (2017).

16. Wen, B. & Noble, W. S. A multi-species benchmark for training and validating mass spectrometry proteomics machine learning models. Scientific Data 11, 1207 (2024).

17. Yilmaz, M., Fondrie, W. E., Bittremieux, W., Oh, S. & Noble, W. S. De novo mass spectrometry peptide sequencing with a transformer model. In Proceedings of the 39th International Conference on Machine Learning, vol. 162, 25514–25522 (2022).

18. Jin, Z. et al. ContraNovo: a contrastive learning approach to enhance de novo peptide sequencing. In Proceedings of the AAAI Conference on Artificial Intelligence (2024).

19. Gessulat, S. et al. Prosit: proteome-wide prediction of peptide tandem mass spectra by deep learning. Nature Methods 16, 509–518 (2019).

20. Brodbelt, J. S. Ion activation methods for peptides and proteins. Analytical Chemistry 88, 30–51 (2016).

21. Aebersold, R. et al. How many human proteoforms are there? Nature Chemical Biology 14, 206–214 (2018).

22. Sun, K. Y. et al. A deep catalogue of protein-coding variation in 983,578 individuals. Nature 631, 583–592 (2024).

23. Sharma, Y. et al. A pan-cancer analysis of synonymous mutations. Nature Communications 10, 2569 (2019).

24. Bittremieux, W. et al. Deep learning methods for de novo peptide sequencing. Mass Spectrometry Reviews 45, 507–526 (2026).

25. Zolg, D. P. et al. Building ProteomeTools based on a complete synthetic human proteome. Nature Methods 14, 259–262 (2017).

26. Rojas Ramírez, C., Yu, F., Polasky, D. A. & Nesvizhskii, A. I. A new detailed mass offset search in MSFragger for improved interpretation of complex PTMs. Journal of Proteome Research 25, 842–851 (2026).

27. Navarro-Jiménez, M. et al. KRAS-targeted therapies in colorectal cancer: a systematic analysis of mutations, inhibitors, and clinical trials. npj Precision Oncology 9, 380 (2025).

28. Zhang, B. et al. Proteogenomic characterization of human colon and rectal cancer. Nature 513, 382–387 (2014).

29. Lundgren, T. J., Clark, P. L. & Champion, M. M. Fit for purpose approach to evaluate detection of amino acid substitutions in shotgun proteomics. Journal of Proteome Research 23, 1263–1271 (2024).

30. Shevchenko, A., Valcu, C.-M. & Junqueira, M. Tools for exploring the proteomosphere. Journal of Proteomics 72, 137–144 (2009).

31. Müller, J. B. et al. The proteome landscape of the kingdoms of life. Nature 582, 592–596 (2020).

32. Abele, M. et al. Proteomic diversity in bacteria: insights and implications for bacterial identification. Molecular & Cellular Proteomics 24, 100917 (2025).

33. Lu, R.-M. et al. Development of therapeutic antibodies for the treatment of diseases. Journal of Biomedical Science 27, 1 (2020).

34. Wang, Y. et al. Simultaneous monitoring of oxidation, deamidation, isomerization, and glycosylation of monoclonal antibodies by liquid chromatography-mass spectrometry method with ultrafast tryptic digestion. mAbs 8, 1477–1486 (2016).

35. Beck, A., Wagner-Rousset, E., Ayoub, D., Van Dorsselaer, A. & Sanglier-Cianférani, S. Characterization of therapeutic antibodies and related products. Analytical Chemistry 85, 715–736 (2013).

36. Bradbury, A. & Plückthun, A. Reproducibility: standardize antibodies used in research. Nature 518, 27–29 (2015).

37. Bandeira, N., Pham, V., Pevzner, P., Arnott, D. & Lill, J. R. Automated de novo protein sequencing of monoclonal antibodies. Nature Biotechnology 26, 1336–1338 (2008).

38. Peng, W., Pronker, M. F. & Snijder, J. Mass spectrometry-based de novo sequencing of monoclonal antibodies using multiple proteases and a dual fragmentation scheme. Journal of Proteome Research 20, 3559–3566 (2021).

39. Beslic, D., Tscheuschner, G., Renard, B. Y., Weller, M. G. & Muth, T. Comprehensive evaluation of peptide de novo sequencing tools for monoclonal antibody assembly. Briefings in Bioinformatics 24, bbac542 (2023).

40. Tran, N. H., Rahman, M. Z., He, L., Xin, L., Shan, B. & Li, M. Complete de novo assembly of monoclonal antibody sequences. Scientific Reports 6, 31730 (2016).

41. Tran, N. H., Zhang, X., Xin, L., Shan, B. & Li, M. De novo peptide sequencing by deep learning. Proceedings of the National Academy of Sciences 114, 8247–8252 (2017).

42. Eloff, K. et al. InstaNovo enables diffusion-powered de novo peptide sequencing in large-scale proteomics experiments. Nature Machine Intelligence 7, 565–579 (2025).

43. Ma, B. & Johnson, R. De novo sequencing and homology searching. Molecular & Cellular Proteomics 11, O111.014902 (2012).

44. Wilmes, P., Heintz-Buschart, A. & Bond, P. L. A decade of metaproteomics: where we stand and what the future holds. Proteomics 15, 3409–3417 (2015).

45. Rodriguez Palomo, I., et al. Benchmarking the identification of a single degraded protein to explore optimal search strategies for ancient proteins. Peer Community Journal 4, e107 (2024).

46. Klaproth-Andrade, D. et al. Modanovo: a unified model for post-translational modification-aware de novo sequencing using experimental spectra from in vivo and synthetic peptides. Molecular & Cellular Proteomics 25, 101501 (2026).

47. Zhang, X. et al. π-PrimeNovo: an accurate and efficient non-autoregressive deep learning model for de novo peptide sequencing. Nature Communications 16, 267 (2025).

48. Zhang, X. et al. Bidirectional representations augmented autoregressive biological sequence generation: application in de novo peptide sequencing. In Advances in Neural Information Processing Systems (2025).

49. Tran, N. H. et al. NovoBoard: a comprehensive framework for evaluating the false discovery rate and accuracy of de novo peptide sequencing. Molecular & Cellular Proteomics 23, 100849 (2024).

50. Mabona, A., et al. De novo peptide sequencing rescoring and FDR estimation with Winnow. Preprint at arXiv 10.48550/arXiv.2509.24952 (2025).

51. 51. Liang, Z. et al. False discovery rate control for trustworthy AI-based de novo peptide sequencing. Preprint at *bioRxiv* 10.64898/2026.06.29.735174 (2026).

52. Fieldhouse, R. This award-winning microscopy image used AI — igniting controversy in a prestigious competition. Nature 10.1038/d41586-026-03086-z (2026).

53. Wang, M. et al. Assembling the community-scale discoverable human proteome. Cell Systems 7, 412–421 (2018).

54. Beslic, D., Tscheuschner, G., Weller, M. G., Renard, B. Y. & Muth, T. Supplementary data for “comprehensive evaluation of peptide de novo sequencing tools for monoclonal antibody assembly” (2022).

55. Giudicelli, V., Chaume, D. & Lefranc, M.-P. IMGT/GENE-DB: a comprehensive database for human and mouse immunoglobulin and T cell receptor genes. Nucleic Acids Research 33, D256–D261 (2005).

56. Loshchilov, I. & Hutter, F. Decoupled weight decay regularization. In International Conference on Learning Representations (2019).

57. Lautenbacher, L. et al. Koina: Democratizing machine learning for proteomics research. Nature Communications 16, 9933 (2025).

58. npysearch contributors. npysearch: BLAST-like biological sequence search for Python. https://github.com/jtruss/npysearch. GitHub repository.

59. Paszke, A. et al. PyTorch: an imperative style, high-performance deep learning library. In Advances in Neural Information Processing Systems, vol. 32 (2019).

60. Karpathy, A. nanogpt. https://github.com/karpathy/nanoGPT (2022). GitHub repository.

61. Levitsky, L. I., Klein, J. A., Ivanov, M. V. & Gorshkov, M. V. Pyteomics 4.0: five years of development of a Python proteomics framework. Journal of Proteome Research 18, 709–714 (2019).

62. Goloborodko, A. A., Levitsky, L. I., Ivanov, M. V. & Gorshkov, M. V. Pyteomics—a Python framework for exploratory data analysis and rapid software prototyping in proteomics. Journal of the American Society for Mass Spectrometry 24, 301–304 (2013).

63. Bittremieux, W. spectrum_utils: a Python package for mass spectrometry data processing and visualization. Analytical Chemistry 92, 659–661 (2020).

64. Cock, P. J. A. et al. Biopython: freely available Python tools for computational molecular biology and bioinformatics. Bioinformatics 25, 1422–1423 (2009).

65. Harris, C. R. et al. Array programming with NumPy. Nature 585, 357–362 (2020).

66. McKinney, W. Data structures for statistical computing in Python. In Proceedings of the 9th Python in Science Conference, 56–61 (2010).

67. Virtanen, P. et al. SciPy 1.0: fundamental algorithms for scientific computing in Python. Nature Methods 17, 261–272 (2020).

68. Pedregosa, F. et al. Scikit-learn: machine learning in Python. Journal of Machine Learning Research 12, 2825–2830 (2011).

69. Hunter, J. D. Matplotlib: a 2D graphics environment. Computing in Science & Engineering 9, 90–95 (2007).

70. Tretyakov, K. matplotlib-venn: area-weighted Venn diagrams for Matplotlib. https://github.com/konstantint/matplotlib-venn. GitHub repository.

71. da Costa-Luis, C. O. et al. tqdm: a fast, extensible progress bar for Python and the CLI. https://github.com/tqdm/tqdm. GitHub repository.

